# Alendronate Conjugate for Targeted Delivery to Bone-Forming Prostate Cancer

**DOI:** 10.1101/2022.09.26.508175

**Authors:** Jossana A. Damasco, Guoyu Yu, Ajay Kumar, Joy Perez, Rio Carlo M. Lirag, Elizabeth M. Whitley, Sue-Hwa Lin, Marites P. Melancon

**Affiliations:** Department of Interventional Radiology, The University of Texas MD Anderson Cancer Center, Houston, TX 77030, USA; Department of Translational Molecular Pathology, The University of Texas MD Anderson Cancer Center, Houston, TX 77030, USA; Department of Biomedical Engineering, Rice University, Houston, TX 77004, USA; Department of Chemistry, Physics, and Engineering, Cameron University–Duncan, Duncan, OK 73533, USA; Department of Veterinary Medicine and Surgery, The University of Texas MD Anderson Cancer Center, Houston, TX 77030, USA; Department of Genitourinary Medical Oncology, The University of Texas MD Anderson Cancer Center, Houston, TX 77030, USA; The University of Texas MD Anderson Cancer Center UT Health Houston Graduate School of Biomedical Sciences, Houston, TX 77030, USA

**Keywords:** alendronate, prostate cancer, bone, near infrared, optical imaging

## Abstract

Bone is the primary metastasis site for lethal prostate cancer, often resulting in poor prognosis, crippling pain, and diminished functioning that drastically reduce both quality of life and survivability. Uniquely, prostate cancer bone metastasis induces aberrant bone overgrowth, due to an increase of osteoblasts induced by tumor-secreted bone morphogenetic protein 4 (BMP4). Conjugating drugs to substances that target the tumor-induced bone area within the metastatic tumor foci would be a promising strategy for drug delivery. To develop such a strategy, we conjugated a near infrared (NIR) fluorescent probe, the dye Cy5.5, to serve as a surrogate for drugs, with alendronate, which targets bone. Characterization, such as infrared spectroscopy, confirmed the synthesis of the Cy5.5-ALN conjugate. The maximum absorbance of free Cy5.5, which was at 675 nm, did not change upon conjugation. Alendronate targeted the bone component hydroxyapatite in a dose-dependent manner up to 2.5 μM, with a maximum of 85% of Cy5.5-ALN bound to hydroxyapatite, while free Cy5.5 alone had 6% binding. In *in vitro* cell binding studies, Cy5.5-ALN bound specifically with mineralized bone matrix of differentiated MC3T3-E1 cells or 2H11 endothelial cells that were induced to become osteoblasts through endothelial-to-osteoblast transition, the underlying mechanism of prostate-cancer-induced bone formation. Neither Cy5.5-ALN nor free Cy5.5 bound to undifferentiated MC3T3-E1 or 2H11 cells. Bone-targeting efficiency studies in non-tumor-bearing mice revealed accumulation over time in the spine, jaw, knees, and paws injected with Cy5.5-ALN, and quantification showed higher accumulation in femurs than in muscle at up to 28 days, while the free Cy5.5 dye was observed circulating without preferential accumulation and decreased over time. There was a linear relationship with fluorescence when the injected concentration of Cy5.5-ALN was between 0.313 and 1.25 nmol/27 g of mouse, as quantified in mouse femurs both *in vivo* and *ex vivo. Ex vivo* evaluation of bone-targeting efficiency in nude mice was 3 times higher for bone-forming C4-2b-BMP4 tumors compared to non-bone-forming C4-2b tumors (p-value < 0.001). Fluorescence microscopy imaging of the tumors showed that Cy5.5-ALN co-localized with the bone matrix surrounding tumor-induced bone, but not with the viable tumor cells. Together, these results suggest that a drug-ALN conjugate is a promising approach for targeted delivery of drug to the tumor-induced bone area in the metastatic foci of prostate cancer.

## 1. Introduction

Prostate cancer is the second-leading cause of cancer deaths among men in the United States, with prognosis primarily determined by whether metastasis develops. One of the hallmarks of prostate cancer bone metastasis is the osteoblastic lesions induced by the tumor [1]. Prostate cancer cells secrete factors, such as bone morphogenetic protein 4 (BMP4), that induce reprogramming of tumor-associated endothelial cells into osteoblasts, which results in bone-forming lesions [2]. This induced aberrant bone formation promotes tumor growth and resistance to therapies [3, 4]. Thus, the ability to selectively target prostate cancer bone metastases could lead to better treatment.

Hydroxyapatite is the major component of bone and binds strongly with bisphosphonates. Bisphosphonates, drugs characterized by phosphorus-carbon-phosphorus (P-C-P) bonds, are routinely used clinically for the prevention and treatment of skeletal-related events. Bisphosphonates work by inhibiting osteoclast bone resorption activity; their varied P-C-P structures allow many different variations with unique physicochemical and biological characteristics. Aside from therapeutic application, bisphosphonates have also been used as targeting ligands to highlight calcification in tumors and vessels using various imaging techniques, such as nuclear imaging [5, 6] and optical imaging in the near infrared (NIR) spectra [7–11]. NIR is desirable because light can penetrate deeper into the tissue since water and naturally occurring fluorochromes have the lowest absorption in this region. The demonstration of specific targeting of bone with the bisphosphonate alendronate in molecular imaging strategies [12] motivated us to investigate the effectiveness of this type of probe for targeting bone-forming prostate cancer bone metastasis in animal models.

In this study, we used an NIR fluorescent probe, the dye Cy5.5, as a surrogate for therapeutic agents that can be attached to the bone-targeting ligand alendronate. We hypothesized that this drug-alendronate conjugate can take advantage of alendronate targeting, resulting in enhanced drug accumulation within the bone-forming prostate cancer metastasis. We show the utility of alendronate conjugated with Cy5.5 NIR dye for targeting tumor-induced bone in an osteogenic prostate tumor model. This probe was utilized to non-invasively track calcification of tumors induced through the presence of BMP4. The probe could be replaced with a small molecular weight drug for specific targeting of osteoblastic prostate cancer tumors.

## 2. Experimental Section

### 2.1. Materials

Sodium alendronate trihydrate (ALN) was obtained from Sigma-Aldrich (St. Louis, MO, USA). Sulfo-Cy5.5-NHS ester (potassium 3-(6-((2,5-dioxopyrrolidin-1-yl)oxy)-6-oxohexyl)-1,1-dimethyl-2-((1E,3E,5E)-5-(1,1,3-trimethyl-6,8-disulfonato-1,3-dihydro-2H-benzo[e]indol-2-ylidene)penta-1,3-dien-1-yl)-1H-benzo[e]indol-3-ium-6,8-disulfonate) was obtained from Lumiprobe (Hunt Valley, MD, USA).

### 2.2. Synthesis of dye-alendronate conjugate

The primary amine of alendronate was conjugated to sulfo-Cy5.5-NHS ester to form a stable amide bond. Specifically, a 5-mM solution of alendronate in H_2_O adjusted to pH = 8-9 using 10 mM sodium hydroxide was added to sulfo-Cy5.5-NHS ester at a 1:2 ALN:Cy5.5 (μmol:μmol) ratio. The reaction mixture was incubated in the dark at room temperature with gentle shaking for about 24 h. The final product, termed “Cy5.5-ALN,” was purified by column chromatography using 65% acetonitrile in water.

### 2.3. Characterization of alendronate-dye conjugate

Cy5.5-ALN was characterized in terms of absorbance in the NIR region using a plate reader (Cytation 5, BioTek, Winooski, VT, USA). Yield was calculated using ε_675nm_=211000 M/cm. The infrared absorption spectra measured using iS5-Nicolet Fourier--transform infrared spectrometer (Thermo Fisher Scientific, Waltham, MA, USA) with standard KBr configuration, equipped with deuterated triglycine sulfate detector, and a single bounce, 45° angle of incidence, AR-coated diamond ATR accessory (iD7 ATR module, Thermo Fisher Scientific, Waltham, MA, USA). For each spectrum, an average of 32 scans was collected at a rate of 0.67 scans/s.

### 2.4. Hydroxyapatite binding affinity

A hydroxyapatite binding assay was used to determine the *in vitro* affinity of the Cy5.5 dye-ALN conjugate for bone matrix. Cy5.5-ALN and Cy5.5 were incubated with hydroxyapatite (10 mg/mL) suspensions with gentle shaking for 1 min at room temperature. The dye-conjugates that bound to hydroxyapatite were separated from unbound dye-conjugates by centrifugation at 1000*g* for 1 min. The absorbance levels of the supernatants were measured using a plate reader (Cytation 5; absorbance 675 nm). The percentage of unbound material was quantified by dividing the final absorbance by the initial absorbance and multiplying by 100. Percentage of binding was determined as the amount unbound subtracted from 100.

### 2.5. *In vitro* binding with MC3T3-E1 and 2H11 cell lines

MC3T3-E1 and 2H11 cells were were purchased from American Type Culture Collection (ATCC, Manasas, VA, USA). MC3T3-E1 cells were cultured in osteoblast-differentiation medium (ODM) containing ascorbic acid and β-glycerol phosphate for 16 to 24 days. 2H11 cells were transitioned into osteoblasts as follows: 2H11 cells were cultured in serum-free Dulbecco’s Modified Eagle Medium (DMEM) overnight followed by treatment with BMP4 (100 ng/ml) for 2 to 3 days. 2H11 cells were then differentiated into osteoblasts by culturing in osteogenic medium (ODM) for 7 to 21 days. Alizarin red staining was used to confirm the mineralization of the osteoblasts; formalin-fixed cells were covered with freshly prepared Alizarin Red S (Sigma Aldrich) solution and incubated at room temperature in the dark for 45 min. For Cy5.5 staining of the control and differentiated cells, the cells were incubated in media containing 6.26 μM Cy5.5 or Cy5.5-ALN for 30 min. The cells were then washed three times with PBS to remove unbound dye prior to fixing with 10% formalin. The fixed cells were then permeabilized with 0.1% Triton X-100 in PBS for 10 min before staining with DAPI. Cellular imaging was then performed using a Nikon ECLIPSE Ts2 (Nikon Instruments Inc., Melville, NY, USA) fluorescence microscope with a Cy filter set (620/50 nm excitation filter, 690/50 nm barrier filter). Image acquisition was performed using NIS Elements v. 5.1101.0.0 (Nikon Instruments Inc., Melville, NY, USA), and the images were analyzed with ImageJ software, v. 1.53q (National Institutes of Health).

### 2.6. Generation of BMP4 overexpressed C4-2b cell line

As we previously described [2], luciferase-expressing C4-2b-BMP4 cells were generated by transfecting luciferase- and Tomato-labeled C4-2b cells (C4-2b-LT) with retrovirus containing cDNAs encoding human BMP4 in the bicistronic retroviral vector pBMN-I-Neo. The cells were then selected by G418 (Corning Inc., Corning, NY, USA). C4-2b-LT-BMP4 cells were maintained in RPMI-1640 medium (Corning) supplemented with 10% fetal bovine serum (Sigma-Aldrich) and 1% penicillin-streptomycin solution (Corning). Cultures were grown in 75-cm^3^ or 175-cm^3^ Nunc EasyYFlask cell culture flasks (Thermo Fisher Scientific) and kept at 37°C inside a Heracell 5% CO_2_ incubator (Thermo Fisher Scientific). Cultures were maintained at 60-80% confluence with media replaced two to three times per week.

### 2.7. Tumor model

All experimentation involving animals was approved by MD Anderson Cancer Center’s Institutional Animal Care and Use Committee. Animals were maintained in facilities approved by AAALAC International and in accordance with current US Department of Agriculture, Department of Health and Human Services, and National Institutes of Health regulations and standards.

As we previously described [13], a bone-forming subcutaneous prostate cancer tumor model was developed using 6- to 8-week-old athymic nude mice (Department of Experimental Radiation Oncology, MD Anderson). Luciferase-expressing C4-2b-LT-BMP4 cells in phosphate-buffered saline (PBS) at a density of 10,000 cells/μL were mixed with Matrigel (Corning) at a 1:1 ratio and kept cold on ice. Sedated mice were subcutaneously inoculated on the right and left upper dorsa with 100 μL of cell suspension containing 1 million cells. Tumor size was measured weekly using bioluminescence imaging. Ectopic bone formation in tumors was monitored using X-ray (Faxitron, Tucson, AR, USA) and micro-computed tomography imaging (GE Healthcare, Wauwatosa, WI, USA).

### 2.8. *In vivo* optical imaging and biodistribution studies

The bone targeting of Cy5.5-ALN in comparison with non-targeted Cy5.5 was evaluated in nude mice. Mice were randomly allocated into two groups (n = 5 mice/group). Prior to pre-injection fluorescence imaging, the mice were anesthetized with 2% isoflurane gas in oxygen. During imaging, the mice were maintained in an anesthetized state with 0.5–1.5% isoflurane in oxygen. Fluorescence optical imaging was done using an IVIS imaging system (200 series) (Xenogen Corp., Alameda, CA, USA) with Cy5.5 filter sets (excitation band/emission band: 615–665/695–770 nm). The field of view was 13.1 cm in diameter. The fluency rate for the NIR fluorescence excitation light was 2 mW/cm^2^. The camera settings included maximum gain, 2 × 2 binning, 640 × 480-pixel resolution, and an exposure time of 0.5 s. After the pre-dye optical imaging, mice were injected retro-orbitally with 5.3 nmol/mouse, 100 μL of Cy5.5 or Cy5.5-ALN. Mice were imaged at t = 1, 3, 5, 9, 16, 21, and 28 days after injection. Relative fluorescence was quantified using Aura Imaging Software v. 3.2 (Spectral Instruments Imaging, Tucson, AZ, USA) or Living Image 4.7.3 software (Perkin Elmer, Waltham, MA, USA).

In a separate experiment, to determine the *in vivo* and *ex vivo* fluorescence response with respect to increasing concentrations of Cy-ALN at different time points, nude mice (n=5/group) were injected retro-orbitally with Cy-ALN with the dose range of 11.5 to 185 nmol/kg (or 0.313 nmol to 5 nmol/27 g mice) in a total volume of 100 μL. The mice were imaged using the IVIS system described above at t = 0, 2, and 96 h. At 96 h after injection, mice were euthanized and the femurs were collected, imaged, and quantified as described previously [13].

Additionally, 20 mice bearing luciferase transfected C4-2b-BMP4 or C4-2b-LT (no BMP4) tumors at an average tumor diameter of 5 mm were injected retro-orbitally with Cy5.5-ALN (targeted) or Cy5.5 (non-targeted) (5 mice/group). Mice were imaged at t = 24 h using the imaging parameters described above. After imaging, mice were euthanized, and major tissues/organs, such as the liver, spleen, kidneys, muscle, gut, heart, lungs, femur, and tumors, were collected for analysis. These tissues/organs were imaged as described above. Sections of tumors were H&E and von Kossa stained and scanned using an Aperio AT2 digital whole-slide scanning system (Leica Biosystems, Buffalo Grove, IL, USA). Fluorescence was observed under the fluorescence microscope as described above.

### 2.9. Statistical analysis

Where applicable, data are presented as mean ± standard deviation and were compared and analyzed using a one-way analysis of variance (ANOVA) test and/or two-tailed Student *t* test, assuming that the data followed a normal distribution. A p value of <0.05 was used to determine statistically significant differences.

## 3. Results and Discussion

### 3.1. Synthesis and characterization of alendronate conjugated to Cy5.5 dye

Cy5.5-ALN was synthesized according to the scheme shown in Fig. 1A. Alendronate, which contains a primary amino group, was attached to the hydroxysuccinimide ester group of sulfonated Cy5.5. Although several strategies for bioconjugation of fluorescent bisphosphonates have been developed [14], we opted for this standard method of conjugation. Purification was done using column chromatography, and the percent yield was 33.16%. Fig. 1B shows the overlaid representative spectra of Cy5.5 alone, alendronate alone, and Cy5.5-ALN, using Fourier transform infrared spectroscopy (FTIR). Broad peaks at 3383 cm^-1^ confirmed the presence of N-H groups, and a peak at 1636 cm^-1^ indicated the C=O of the amide group in the Cy5.5-ALN conjugate. UV-vis absorption showed that Cy5.5-ALN had a maximum absorption at 675 nm, which was the same as that of free Cy5.5 (Fig. 1C). Hydroxyapatite binding was used to evaluate the binding affinity of targeted versus non-targeted dye to bone. Both targeted and non-targeted solutions were blue due to the presence of the dye. When Cy5.5-ALN was incubated with hydroxyapatite, the white hydroxyapatite particles turned blue, while the supernatant became colorless (Fig. 2A). In contrast, when non-targeted Cy5.5 was incubated with hydroxyapatite, the hydroxyapatite particles remained white and the supernatant turned blue, indicating that free Cy5.5 did not bind to hydroxyapatite efficiently and that alendronate leads to binding of Cy5.5-ALN to hydroxyapatite. Quantification of percentage of binding was done at different Cy5.5 concentrations using absorbance at 675 nm; it showed a direct relationship between Cy5.5-ALN concentration and amount of hydroxyapatite binding (Fig. 2B). Furthermore, Cy5.5-ALN bound to hydroxyapatite almost instantaneously, and the binding plateaued and reached a maximum of 85%. In contrast, non-targeted Cy5.5 had very low binding (up to 6%) to hydroxyapatite after incubation (Fig. 2C).

**Fig. 1.**
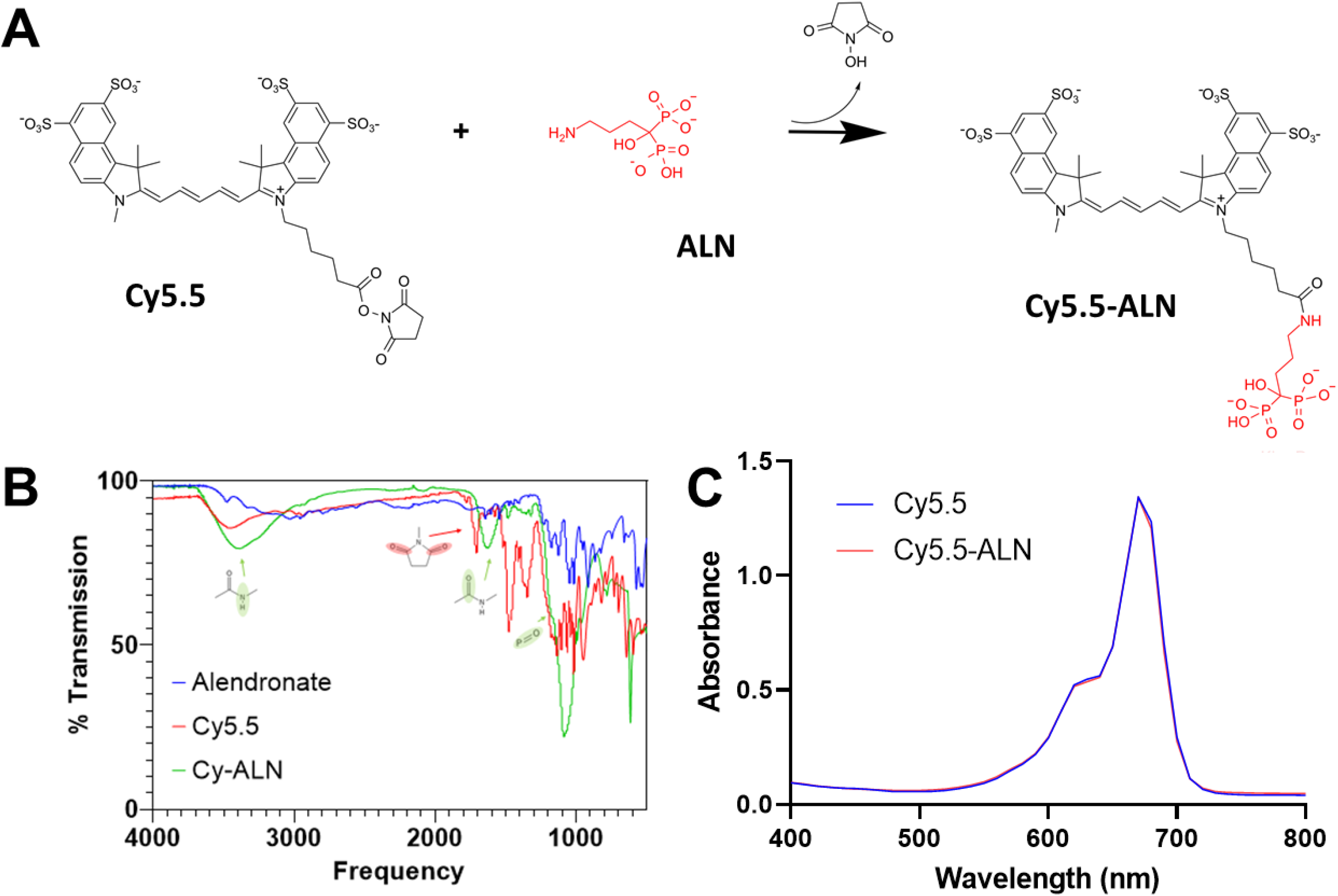
Synthesis and characterization of alendronate-conjugated dye. (A) Scheme for the synthesis of alendronate (ALN)-conjugated Cy5.5 (Cy5.5-ALN). (B) Infrared spectroscopy of alendronate, Cy5.5, and Cy5.5-ALN. (C) Absorbance of Cy5.5 and Cy5.5-ALN in the NIR region.

**Fig. 2.**
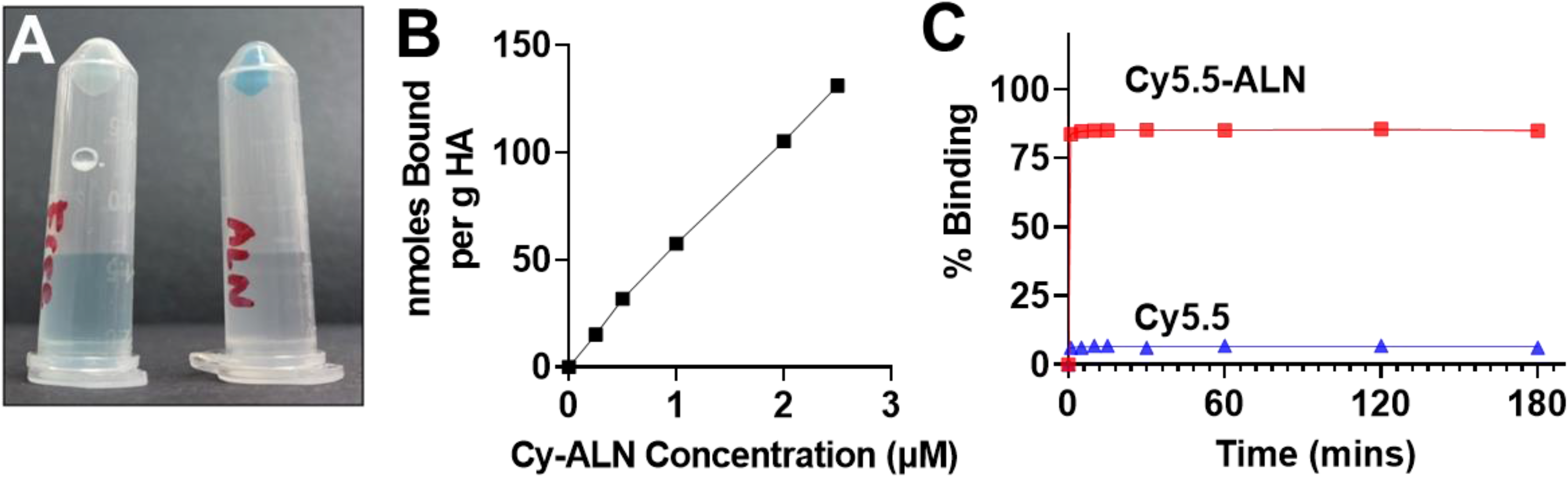
Hydroxyapatite binding. (A) Photograph of binding of Cy5.5 and Cy5.5-ALN to hydroxyapatite. (B) Concentration-dependent binding of Cy5.5-ALN. (C) Binding of Cy5.5-ALN and Cy5.5 to hydroxyapatite over time.

### 3.2. Binding of Cy5.5-ALN to mineralized bone matrix *in vitro*

We next examined the binding of Cy5.5-ALN to mineralized bone matrix in osteoblast culture. We first used MC3T3-E1 cells as our *in vitro* assay system. MC3T3-E1 is a cell line derived from mouse calvaria and, when cultured in ODM (osteogenic medium), differentiate into mature osteoblasts with mineralized bone matrix, as shown in an Alizarin red assay (Fig. 3A). In this study, MC3T3-E1 cells were cultured in ODM for 16 days. MC3T3-E1 cells without differentiation were used as a control. Cy5.5-ALN or Cy5.5 was added to the differentiated MC3T3-E1 cells for 30 min, and the cells were imaged using an inverted fluorescence microscope using the Cy5.5 emission filter. Cy5.5-ALN was found to more selectively bind to differentiated MC3T3-E1 cells than to undifferentiated MC3T3-E1 cells (Fig. 3B). The non-targeted Cy5.5 showed very low binding to either undifferentiated or differentiated MC3T3-E1 cells (Fig. 3B, left panels). These results indicate that Cy5.5-ALN has higher binding activity than Cy5.5 to mineralized bone matrix of MC3T3-E1 cells.

**Fig. 3.**
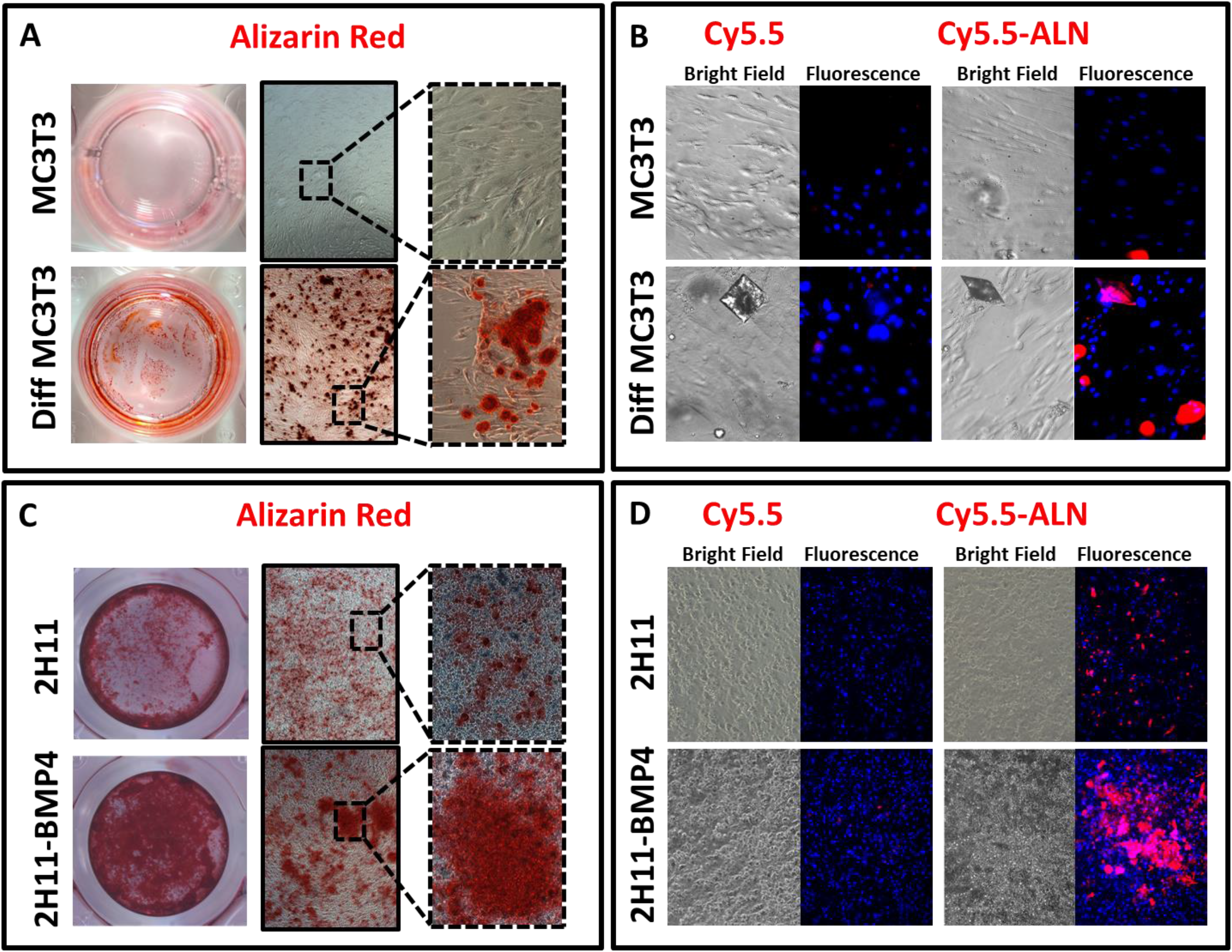
*In vitro* binding of Cy5.5 and Cy5.5-ALN with mineralized versus non-mineralized cells. (A, C) Differentiated MC3T3-E1 (A) and 2H11-BMP4 (C) cells show more Alizarin red staining, indicating more mineralization as compared with undifferentiated cells. (B, D) Fluorescence microscopy images show binding of Cy5.5-ALN to mineralized MC3T3-E1 (B) and 2H11-BMP4 (D) cells (red overlaid with blue DAPI nuclear staining) compared to no or minimal binding with undifferentiated MC3T3-E1 and 2H11 cells. Non-targeted Cy5.5 did not bind with mineralized or non-mineralized cells.

We previously showed that the bone-forming phenotype of prostate cancer bone metastasis is through BMP4-induced endothelial-to-osteoblast (EC-to-OSB) transition [2, 13]. Treatment of 2H11 endothelial cells with BMP4 induced EC-to-OSB transition, leading to formation of mineralized bone matrix [2]. We also examined the binding of Cy5.5-ALN to mineralized bone matrix from EC-to-OSB transition. We treated 2H11 cells, an endothelial cell line, with BMP4 and ODM for 14 days to induce EC-to-OSB transition and osteoblast differentiation, which was confirmed by Alizarin red staining (Fig. 3C). Cy5.5-ALN was found to preferentially bind to the BMP4-treated 2H11 cells compared to the control 2H11 cells (Fig. 3D). Cy5.5 did not exhibit significant binding to either differentiated or undifferentiated 2H11 cells (Fig. 3D). These results further support the preferential binding of Cy5.5-ALN to mineralized bone matrix.

### 3.3. *In vivo* bone targeting of Cy5.5-ALN in non-tumor-bearing mice

To compare the efficacy of Cy5.5-ALN versus Cy5.5 in targeting bones in nude mice, we used optical fluorescence imaging, which is a fast and real-time modality for monitoring the localization of Cy5.5-ALN. Mice were injected retro-orbitally with a single dose of 5 nmol/mouse Cy5.5-ALN (targeting) or Cy5.5 (non-targeting) and imaged at t = 1, 2, 3, 5, 9, 16, 21, and 28 days after injection. NIR fluorescence optical imaging showed high fluorescence for Cy5.5-injected mice at day 1, which slowly decreased until day 9, and no fluorescence was detected from day 16 onwards (Fig. 4A). In comparison, after initial diffused distribution, Cy5.5-ALN had increased accumulation in the skull, spine, tail, and paws (Fig. 4A). Quantification of the fluorescence intensity in radiance showed accumulation of Cy5.5-ALN in the leg increased from 6.3 x10^9^ photons/s at day 1 up to 1.4 x10^10^ photons/s on day 9 and then decreased to 1.2 x10^10^ photons/s on day 28; accumulation in the muscle remained low, 2.8 x 10^9^ photons/s at day 1 to 1.7 x 10^9^ photons/s at day 28 (Fig. 4B). In contrast, non-targeted Cy5.5 had high fluorescence intensity (1.1 x 10^10^ photons/s) in both femurs and muscle on day 1, and both decreased over time until reaching the baseline intensity of 1.1 x 10^9^ photons/s on day 9 (Fig. 4B). A large difference in fluorescence intensity was observed in the femurs between the targeted Cy5.5-ALN and non-targeted Cy5.5 starting from day 1, and thus 96 hours was used as the last time point for the subsequent experiments.

**Fig. 4.**
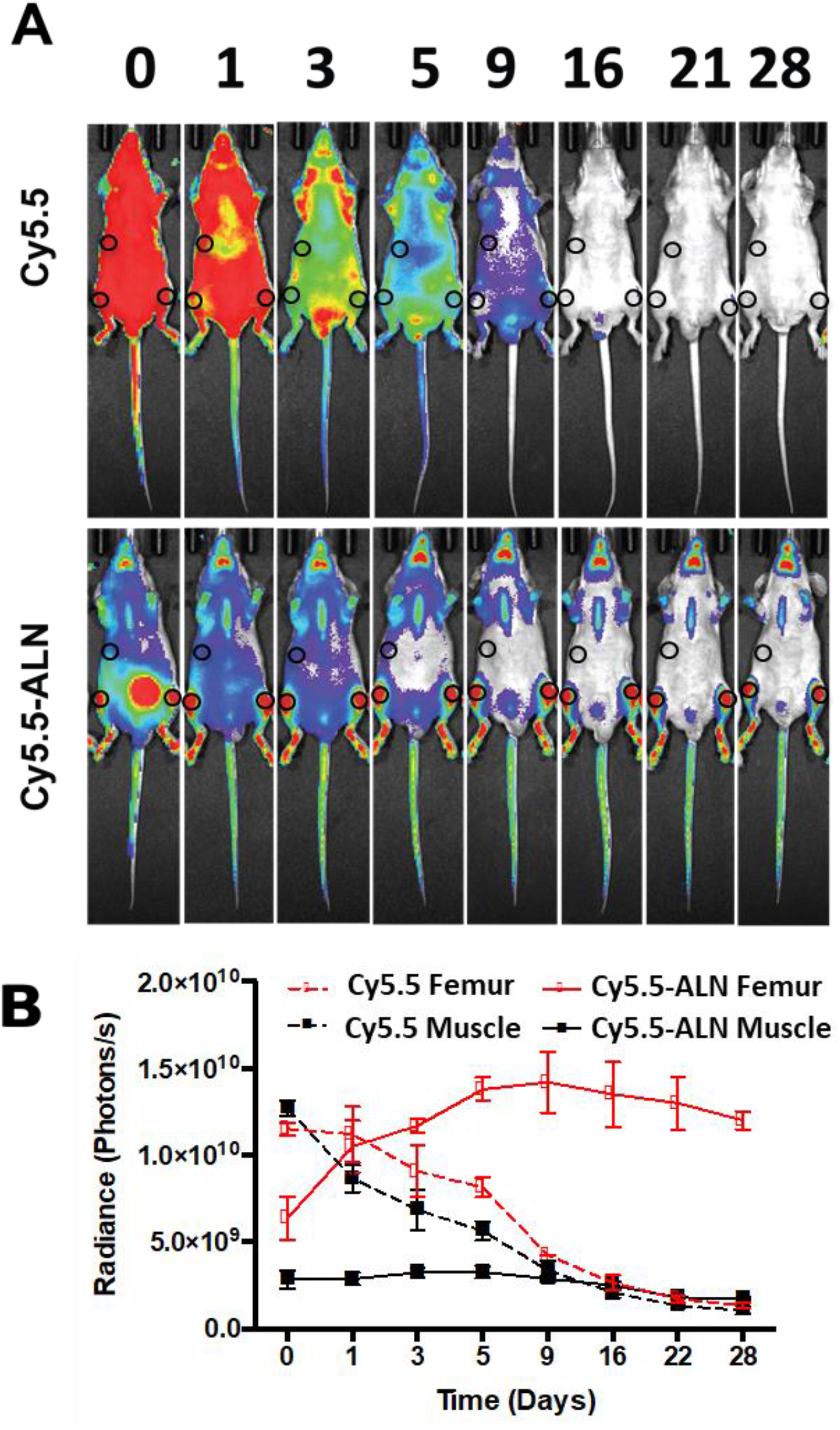
Bone-targeting efficiency of Cy5.5-ALN versus Cy5.5 in mice. (A) NIR optical imaging of nude mice before (t = 0) and after intraorbital injection of Cy5.5-ALN or Cy5.5 at different time points (t = 1, 3, 5, 9, 16, 21, and 28 days). The small circles indicate the regions of interest for muscle (top circle) and leg (bottom circles) taken for quantification. (B) Quantification of fluorescence in photons/s using Aura 3.2 software.

The differential binding of Cy5.5-ALN to bone is likely concentration dependent. We thus determined the concentration dependence of the conjugate’s uptake in mice *in vivo* at different time points. Fig. 5A shows that as the Cy5.5-ALN concentration was increased, the fluorescence in the legs also increased. The maximum increase was observed at 2 h for low concentrations ranging from 0.313 to 1.25 nmol/mouse (Fig. S1). However, the background fluorescence was also very high owing to the circulation of the conjugate, which was almost cleared at 6 h for mice injected with the lowest concentration of Cy5.5-ALN (Fig. S2). Signal intensity in the femur remained high even at 96 h. We then looked at linearity of fluorescence *in vivo* at 2 and 96 h after injection (Fig. 5A). Our results show linear uptake from 0.313 to 1.25 nmol/mouse (R2 = ~0.98), but not at 2.5 to 5 nmol/mouse, at both 2 and 96 h. The same trend was observed *ex vivo* in isolated femurs at 96 h (Fig. 5B), with a linear relationship at the lower concentration (0.313 to 1.25 nmol/mouse) but not at the higher concentration (2.5 to 5 nmol/mouse).

**Fig. 5.**
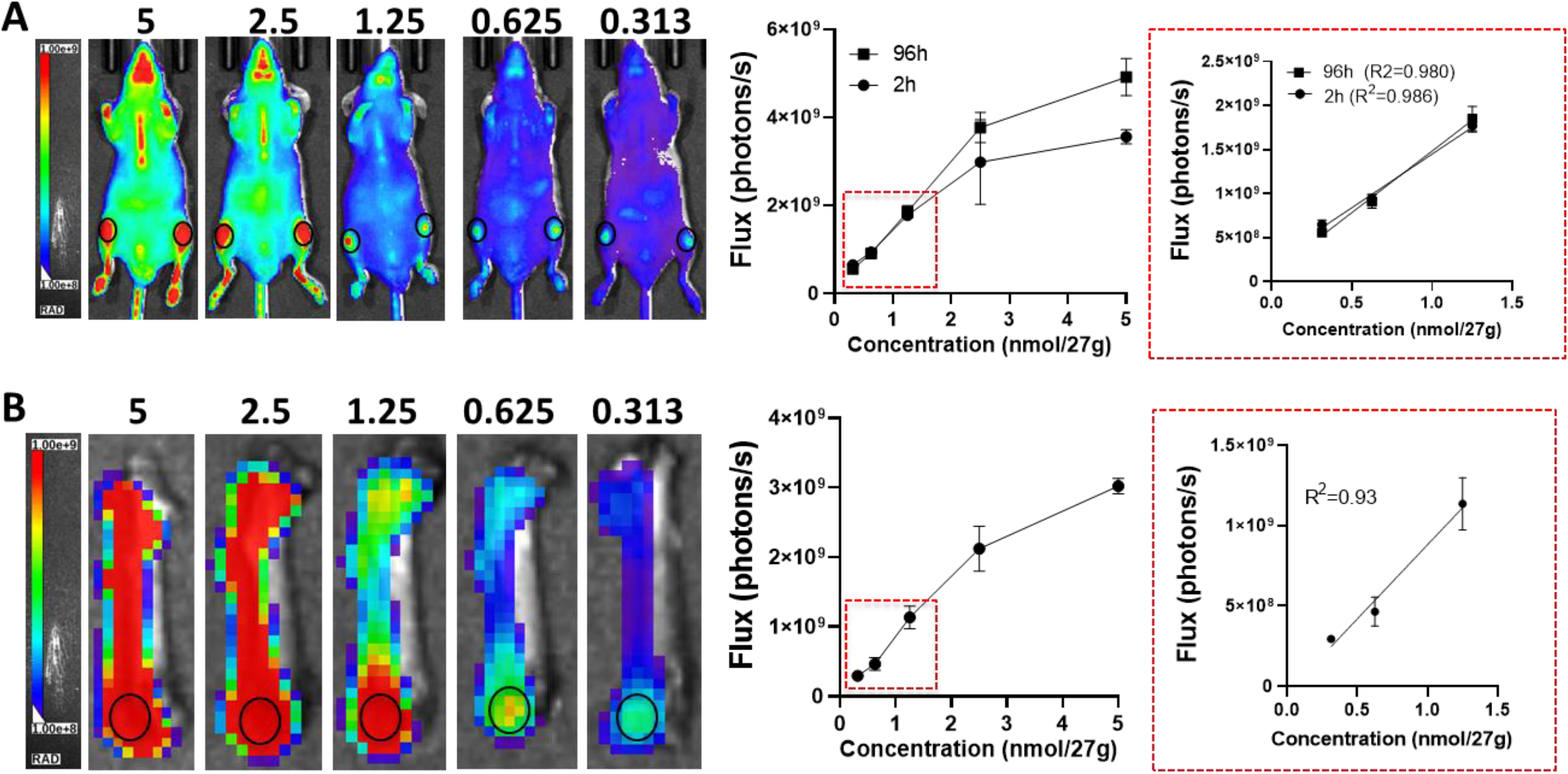
Dose dependence of Cy5.5-ALN uptake in bones. (A) *In vivo* fluorescence imaging of Cy5.5-ALN in nude mice at different concentrations at 2 h. The dose-dependent increase in fluorescence was similar at all time points. (B) *Ex vivo* bone imaging. Femurs isolated at 96 h after injection show a linear relationship between fluorescence and concentration at low (0.313 to 1.25 nmol/mouse) but not high (2.5 to 5 nmol/mouse) concentrations. Quantification was done using Aura 3.2 software.

### 3.4. Targeting of Cy5.5-ALN to prostate-cancer-induced bone *in vivo*

We have shown that alendronate has a high affinity for hydroxyapatite, which is present on the surfaces of bones that are undergoing active remodeling. Because osteogenic prostate tumor contains cancer-induced new bone, we hypothesize that Cy5.5-ALN will selectively target tumor-induced bone, in addition to normal bone. To test this, we inoculated nude mice with bone-forming C4-2b-BMP4 cells and non-bone-forming C4-2b cells. C4-2b-BMP4 cells, established by transfecting C4-2b cells with BMP4 [13], generate ectopic bone in tumor when implanted subcutaneously. C4-2b and C4-2b-BMP4 cells were injected subcutaneously in nude mice. After 4 weeks, mice were injected with either Cy5.5-ALN or Cy5.5, using the lowest concentration, i.e., 0.313 nmol/mouse, and specific targeting was monitored using fluorescence imaging at the NIR range. As C4-2b and C4-2b-BMP4 tumor cells express luciferase, biolumiscence (BLI) was used to locate the tumors. Non-targeted Cy5.5 had non-specific accumulation in the entire mouse in both C4-2b- and C4-2b-BMP4-injected mice (Fig. 6). In contrast, targeted Cy5.5-ALN had increased accumulation in the skull, spine, tail, and paws, as previously observed in non-tumor-bearing mice. Importantly, there was also increased Cy5.5-ALN fluorescence in C4-2b-BMP4 tumors compared to C4-2b tumors (tumor/muscle ratio, 1.45 and 0.88, respectively; p-value = 0.009).

**Fig. 6.**
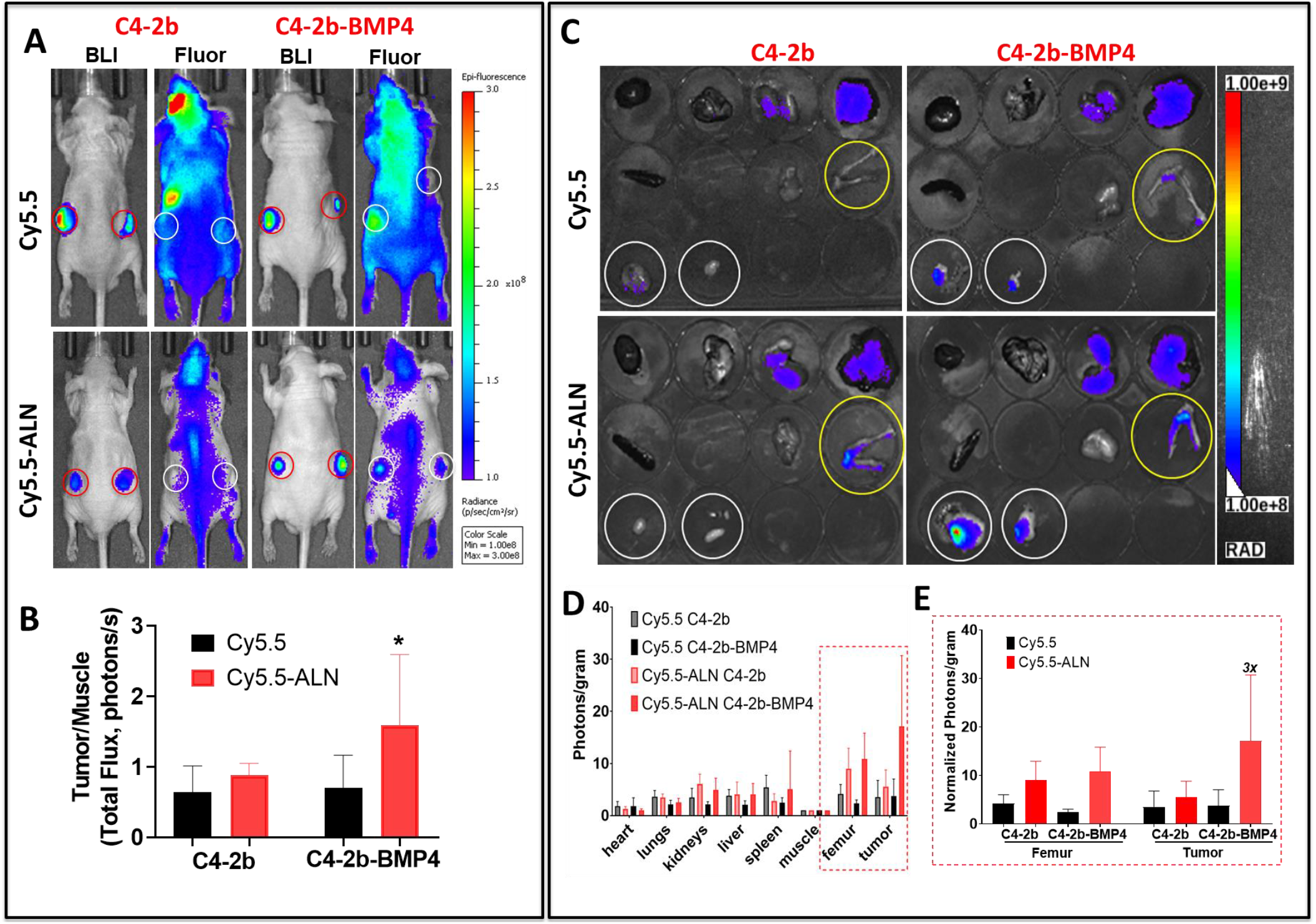
*In vivo* and *ex vivo* analyses of targeted Cy5.5-ALN in mice bearing bone-forming prostate cancer tumors. (A) Bioluminescence (BLI) and fluorescence (Fluor) imaging of nude mice bearing C4-2b or C4-2b-BMP4 tumors at 24 h after intraorbital injection of Cy5.5-ALN or Cy5.5. Red circles indicate areas quantified for bioluminescence, while white circles are areas for fluorescence. (B) Quantification of fluorescence in photons/s using Living Image 4.7.3 software (Perkin Elmer, Waltham, MA, USA). A statistically significant increase in fluorescence in the tumor area was observed with Cy5.5-ALN injected into C4-2b-BMP4 tumor-bearing mice as compared with C4-2b tumor-bearing mice (p-value < 0.001). No significant difference in fluorescence between C4-2b-BMP4 and C4-2b tumors was observed in mice injected with the non-targeted Cy5.5. (C) Fluorescence imaging of the various organs, tumor, and leg at 24 h after injection of Cy5.5-ALN or Cy.5.5 in nude mice bearing C4-2b or C4-2b-BMP4 tumors. Top row: Gut, liver, spleen, kidneys. Middle row: Lungs, heart, muscle, leg (circled in yellow). Bottom row: Tumors (circled in white). (D) Quantification of fluorescence in photons/s using Living Image 4.7.3 software. There was an increase in accumulation of Cy5.5-ALN in the legs of mice bearing C4-2b or C4-2b-BMP4 tumors, respectively, compared with Cy5.5, as expected. Accumulation of Cy5.5-ALN in C4-2b-BMP4 tumors was three times higher than that of non-targeted Cy5.5 (p-value < 0.001).

### 3.5. Biodistribution

Biodistribution studies upon organ dissection at day 1 showed minimal accumulation of the agents in gut, liver, spleen, kidneys, lungs, heart, and muscle (1.8 x 10^9^ – 8.3 x 10^9^ photons/s) in mice bearing C4-2b or C4-2b-BMP4 tumors. In the legs, there was increased accumulation of Cy5.5-ALN in mice bearing either C4-2b-BMP4 (6.7 x 10^9^ photons/s) or C4-2b tumors (4.8 x 10^9^ photons/s) (Fig. 6) compared to Cy5.5 (3.3 x 10^9^ photons/s), as expected. Importantly, accumulation of Cy5.5-ALN in tumors was 3 times higher for the osteogenic C4-2b-BMP4 tumors (1.3 x 10^10^ photons/s) compared to non-osteogenic C4-2b tumors (4.2 x 10^9^ photons/s) (p-value < 0.001). Thus, *in vivo* and *ex vivo* optical imaging revealed significant accumulation of Cy5.5-ALN in C4-2b-BMP4 tumors at 1 day after injection of the dye (Fig. 6).

### 3.6. Histology

Hematoxylin and eosin (H&E) staining of whole tumors revealed ectopic bone formation in C4-2b-BMP4 but not in C4-2b tumors (Fig. 7). Von Kossa staining further confirmed mineralized ectopic bone in C4-2b-BMP4 tumors but not in C4-2b tumors (Fig. 7). Under a fluorescence microscope, we found increased Cy5.5-ALN fluorescence signal in C4-2b-BMP4 tumors compared with C4-2b tumors. Free Cy5.5 without alendronate did not have any fluorescence signal in either C4-2b-BMP4 or C4-2b tumors (Fig. 7). Furthermore, fluorescence imaging of the tumors showed that Cy5.5-ALN co-localized with the bone matrix surrounding tumor-induced bone, but not with the viable tumor cells (Fig. 7B). These results confirm that Cy5.5-ALN selectively binds to the mineralized areas of the tumor.

**Fig. 7.**
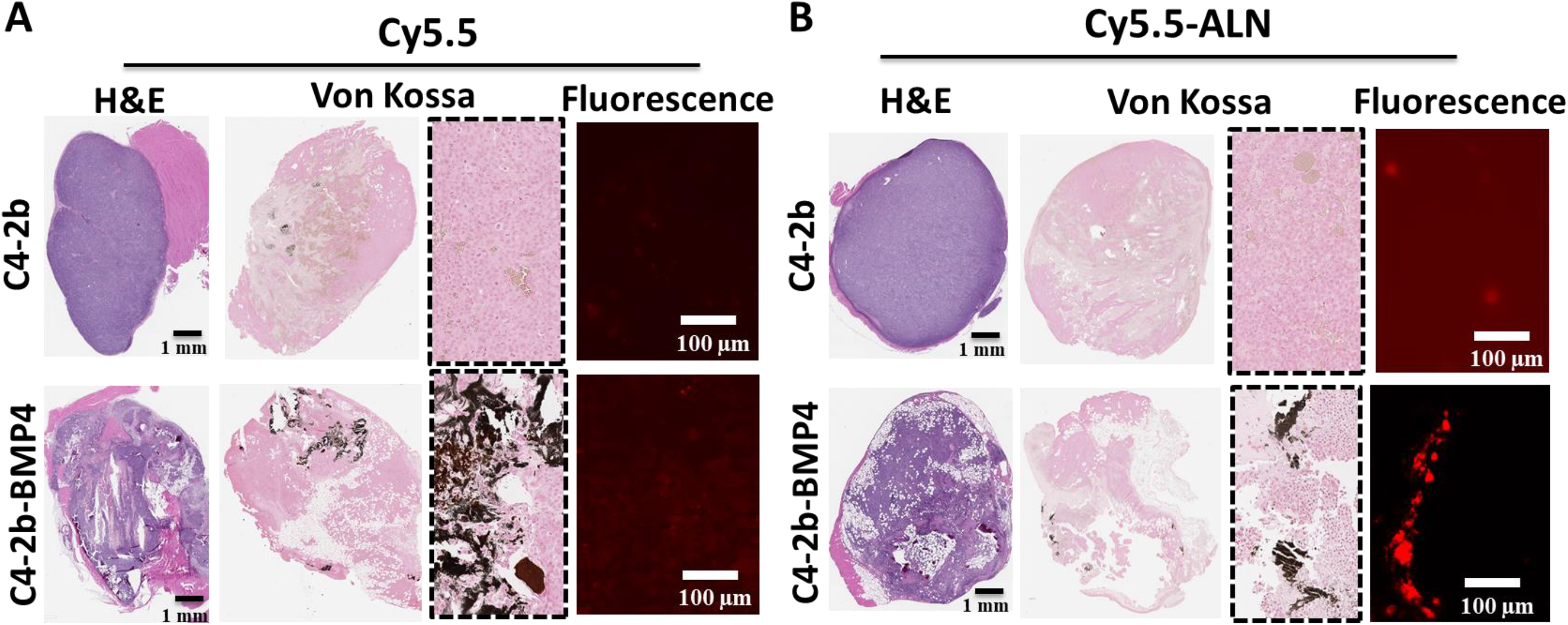
H&E staining, von Kossa staining, and fluorescence of tumors from nude mice bearing C4-2b and C4-2b-BMP4 tumors. (A) Tumor-bearing mice were injected with Cy5.5. (B) Tumor-bearing mice were injected with Cy5.5-ALN.

## 4. Conclusion

We demonstrated that we are able to synthesize Cy5.5-ALN without reducing the ability of alendronate to target hydroxyapatite. *In vitro* and *in vivo* studies showed that Cy5.5-ALN targets mineralized components of bone-forming tumors. These findings demonstrate that a drug-ALN conjugate is a promising approach for targeted delivery of drug to the tumor-induced bone area of bone-metastatic prostate cancer.

## Acknowledgments

This work was supported in part by grants from the National Institutes of Health (NIH)/National Heart, Lung, and Blood Institute [5R01HL141831-03 and 1R01HL159960-01A1 to M.P.M.], NIH/National Cancer Institute [R01CA174798 and 5P50CA140388 to S.-H.L.], and Cancer Prevention and Research Institute of Texas [RP150179 and RP190252 to S.-H.L.]. MD Anderson’s Research Animal Support Facility and Small Animal Imaging Facility are supported by NIH/NCI through MD Anderson’s Cancer Center Support Grant [P30CA016672]. The authors would like to acknowledge Sunita Patterson in MD Anderson’s Research Medical Library for editing the manuscript.

## References

[1] C.J. Logothetis, S.-H. Lin, Osteoblasts in prostate cancer metastasis to bone, Nature Reviews Cancer 5(1) (2005) 21–28.

[2] G. Yu, P. Shen, Y.-C. Lee, J. Pan, J.H. Song, T. Pan, S.-C. Lin, X. Liang, G. Wang, T. Panaretakis, C.J. Logothetis, G.E. Gallick, L.-Y. Yu-Lee, S.-H. Lin, Multiple pathways coordinating reprogramming of endothelial cells into osteoblasts by BMP4, iScience 24(4) (2021) 102388–102388.

[3] Y.C. Lee, C.J. Cheng, M.A. Bilen, J.F. Lu, R.L. Satcher, L.Y. Yu-Lee, G.E. Gallick, S.N. Maity, S.H. Lin, BMP4 promotes prostate tumor growth in bone through osteogenesis, Cancer Res 71(15) (2011) 5194–203.

[4] Y.C. Lee, S.C. Lin, G. Yu, C.J. Cheng, B. Liu, H.C. Liu, D.H. Hawke, N.U. Parikh, A. Varkaris, P. Corn, C. Logothetis, R.L. Satcher, L.Y. Yu-Lee, G.E. Gallick, S.H. Lin, Identification of Bone-Derived Factors Conferring De Novo Therapeutic Resistance in Metastatic Prostate Cancer, Cancer Res 75(22) (2015) 4949–59.

[5] K. Ogawa, T. Mukai, Y. Arano, M. Ono, H. Hanaoka, S. Ishino, K. Hashimoto, H. Nishimura, H. Saji, Development of a Rhenium-186-Labeled MAG3-Conjugated Bisphosphonate for the Palliation of Metastatic Bone Pain Based on the Concept of Bifunctional Radiopharmaceuticals, Bioconjugate Chemistry 16(4) (2005) 751–757.

[6] M.G. Lam, J.M. de Klerk, P.P. van Rijk, B.A. Zonnenberg, Bone seeking radiopharmaceuticals for palliation of pain in cancer patients with osseous metastases, Anticancer Agents Med Chem 7(4) (2007) 381–97.

[7] A. Zaheer, R.E. Lenkinski, A. Mahmood, A.G. Jones, L.C. Cantley, J.V. Frangioni, In vivo near-infrared fluorescence imaging of osteoblastic activity, Nat Biotechnol 19(12) (2001) 1148–54.

[8] R.E. Lenkinski, M. Ahmed, A. Zaheer, J.V. Frangioni, S.N. Goldberg, Near-infrared fluorescence imaging of microcalcification in an animal model of breast cancer, Acad Radiol 10(10) (2003) 1159–64.

[9] K.R. Bhushan, E. Tanaka, J.V. Frangioni, Synthesis of Conjugatable Bisphosphonates for Molecular Imaging of Large Animals, Angewandte Chemie International Edition 46(42) (2007) 7969–7971.

[10] Y. Zilberman, I. Kallai, Y. Gafni, G. Pelled, S. Kossodo, W. Yared, D. Gazit, Fluorescence molecular tomography enables in vivo visualization and quantification of nonunion fracture repair induced by genetically engineered mesenchymal stem cells, Journal of Orthopaedic Research 26(4) (2008) 522–530.

[11] A.M. Sim, N.A. Rashdan, L. Cui, A.J. Moss, F. Nudelman, M.R. Dweck, V.E. MacRae, A.N. Hulme, A novel fluorescein-bisphosphonate based diagnostic tool for the detection of hydroxyapatite in both cell and tissue models, Scientific Reports 8(1) (2018) 17360.

[12] R.A. Nadar, N. Margiotta, M. Iafisco, J. van den Beucken, O.C. Boerman, S.C.G. Leeuwenburgh, Bisphosphonate-Functionalized Imaging Agents, Anti-Tumor Agents and Nanocarriers for Treatment of Bone Cancer, ADVANCED HEALTHCARE MATERIALS 6(8) (2017).

[13] S.-C. Lin, Y.-C. Lee, G. Yu, C.-J. Cheng, X. Zhou, K. Chu, M. Murshed, N.-T. Le, L. Baseler, J.-i. Abe, K. Fujiwara, B. deCrombrugghe, C.J. Logothetis, G.E. Gallick, L.-Y. Yu-Lee, S.N. Maity, S.-H. Lin, Endothelial-to-Osteoblast Conversion Generates Osteoblastic Metastasis of Prostate Cancer, Developmental Cell 41(5) (2017) 467–480.e3.

[14] S. Sun, K.M. Błażewska, A.P. Kadina, B.A. Kashemirov, X. Duan, J.T. Triffitt, J.E. Dunford, R.G.G. Russell, F.H. Ebetino, A.J. Roelofs, F.P. Coxon, M.W. Lundy, C.E. McKenna, Fluorescent Bisphosphonate and Carboxyphosphonate Probes: A Versatile Imaging Toolkit for Applications in Bone Biology and Biomedicine, Bioconjugate chemistry 27(2) (2016) 329–340.

