## Supplemental Figure 1 for "Alendronate Conjugate for Targeted Delivery to Bone-Forming Prostate Cancer"

Houston, TX 77030, USA

[c]Department of Biomedical Engineering

Rice University

Houston, TX 77004, USA

[d]Department of Chemistry, Physics, and Engineering

Cameron University–Duncan

Duncan, OK 73533, USA

[g]The University of Texas MD Anderson Cancer Center UT Health Houston Graduate School of Biomedical Sciences

Houston, TX 77030, USA

[†]Co-first Authors

*Corresponding Authors:

Prof. M. P. Melancon

Department of Interventional Radiology, Unit 1471, The University of Texas MD Anderson Cancer Center, 1515 Holcombe Boulevard, Houston, TX 77030, USA

Prof. S-H Lin

Department of Translational Molecular Pathology, Unit 0089, The University of Texas MD Anderson Cancer Center, 1515 Holcombe Boulevard, Houston, TX 77030, USA

Keywords: *alendronate, prostate cancer, bone, near infrared, optical imaging*

**Supplementary Figures:**

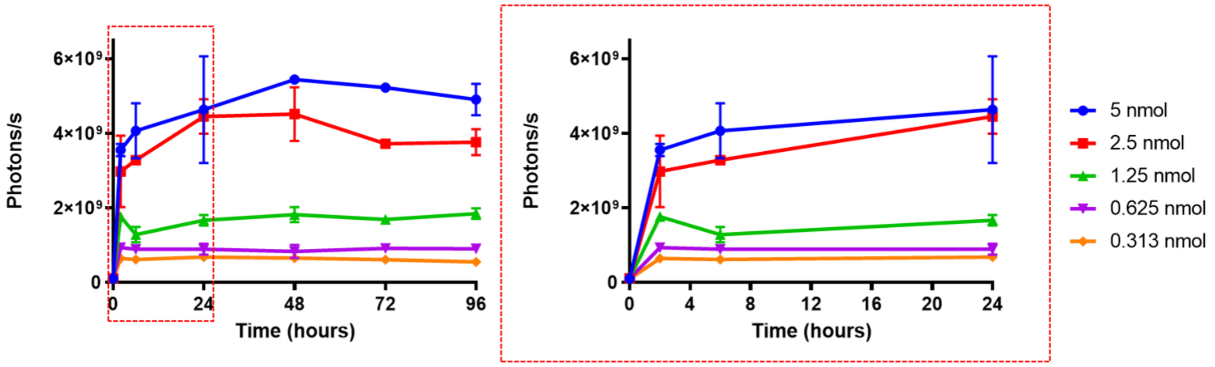


Fig. S1. Left: Dose-dependent uptake of Cy5.5-ALN in non-tumor-bearing mice at all time points (t=0, 2, 6, 24, 48, 72, and 96 h). Right: Zoomed graph showing uptake during the first 24 h.

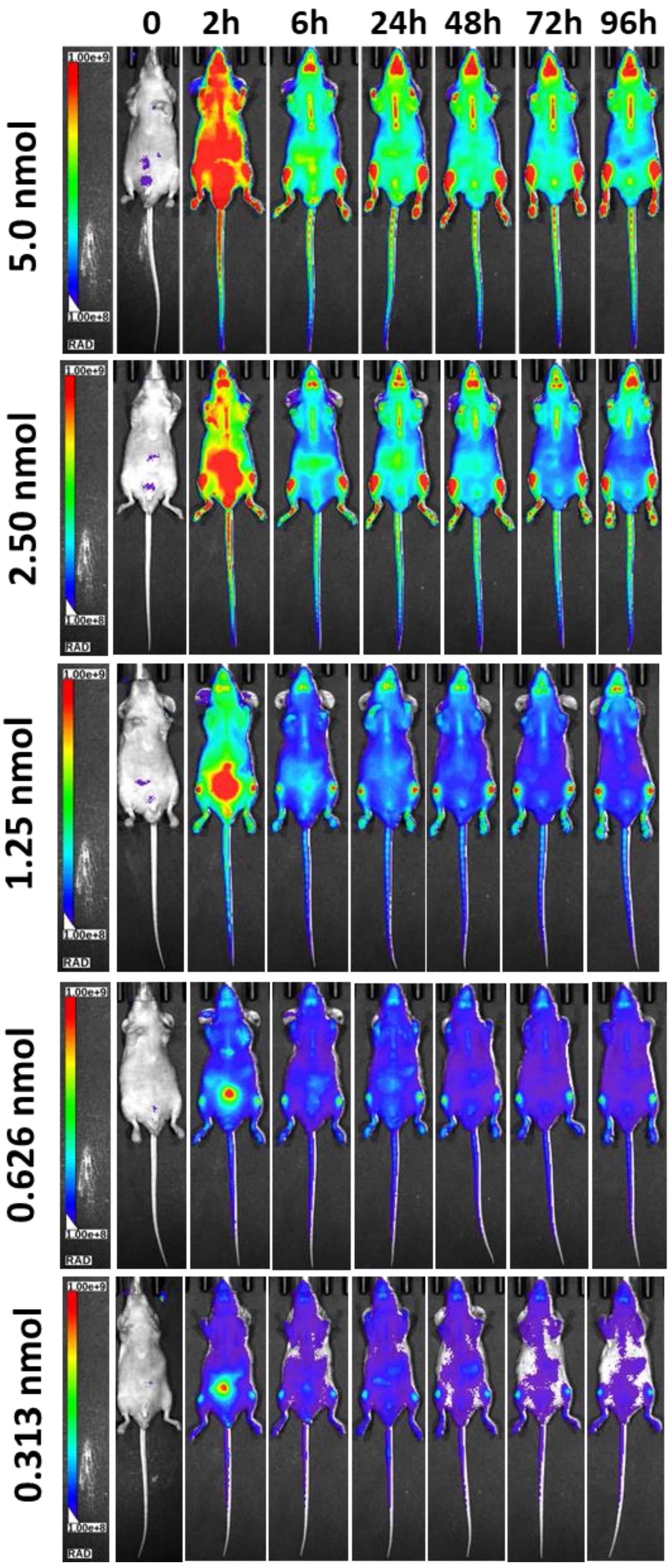


Fig. S2. Representative images of the fluorescence uptake in non-tumor-bearing mice at each time point using different concentrations of Cy5.5-ALN.
